# Detection of Chronic Wasting Disease prions in fetal tissues *of* free-ranging white-tailed deer

**DOI:** 10.1101/2021.03.13.435248

**Authors:** AV Nalls, EE McNulty, A Mayfield, JM Crum, MK Keel, EA Hoover, MG Ruder, CK Mathiason

## Abstract

The transmission of chronic wasting disease (CWD) has largely been attributed to contact with infectious prions shed in excretions (saliva, urine, feces, blood) by direct animal-to-animal exposure or indirect contact with the environment. Less-well studied has been the role mother-to-offspring transmission may play in the facile transmission of CWD. We asked whether such extensive spread may also be due to mother-to-offspring transmission, perhaps before birth. We thereby focused on a population of white-tailed deer from West Virginia, USA, in which CWD has been detected. Fetal tissues, ranging from 113 to 158 days of gestation, were harvested from the uteri of CWD+ dams in the asymptomatic phase of infection. Using serial protein misfolding amplification (sPMCA), we detected evidence of prion seeds in 6 of 14 *in utero* harvested fetuses, with earliest detection at 113 gestational days. This is the first report of CWD detection in free ranging white-tailed deer fetal tissues. Further investigation within cervid populations across North America will help define the role and impact of mother-to-offspring vertical transmission of CWD.

## Introduction

Investigations into the transmission dynamics of chronic wasting disease (CWD) have primarily focused on the presence of infective material in bodily fluids and excretions of infected cervids ^1–4^. The prevailing hypothesis is that contact with infectious prions shed by infected cervids via animal-to-animal contact or presumed ingestion of prions from contaminated environments represent the majority of CWD spread among cervids. Less well-studied has been the role of prion transmission from mother to offspring.

Prion transmission from mother to offspring has been demonstrated for sheep scrapie ^5–7^. The first evidence for maternal prion transmission came from observational studies revealing that the incidence of scrapie infections increased during the lambing season ^6^. Further investigation led to reports of PrP^Scrapie^ deposition within maternal and fetal tissues ^8–15^ and the presence of infectious prions within placental tissues ^16,17^ fetal tissues ^18^, embryos ^14^ and milk of scrapie-infected dams ^19–23^. This effectively demonstrated a role for maternal transmission, with increasing evidence for prebirth fetal exposure to scrapie.

The evidence for prebirth CWD exposure and transmission is also mounting. We previously reported mother-to-offspring transmission and the detection of CWD amyloid seeding activity (prions) in maternal and fetal tissues harvested from experimentally-infected Reeve’s muntjac (*Muntiacus reeves*)^24^. Further studies in our maternal infection model led to the discovery of prion infectivity within the pregnancy microenvironment (uterus, ovary, placentomes) ^25^. This piqued our interest to investigate the biological relevance of mother-of-offspring transmission in free-range cervid populations. These studies resulted in demonstration of prions within *in utero*-derived fetal tissues harvested from naturally exposed free-ranging CWD+ asymptomatic Rocky Mountain elk (*Cervus canadensis*) cows ^26^.

To explore potential differences between cervid species in prebirth dissemination of prions from mother-to-offspring we extended our studies to investigate *in utero*-derived fetal tissues from white-tailed deer (*Odocoileus virginianus*). Here we used serial protein misfolding cyclic amplification (sPMCA) to assess fetal tissues harvested from asymptomatic CWD+ does from West Virginia, USA. We report, for the first time, prion seeding activity within fetal tissues of naturally exposed free-ranging asymptomatic CWD+ white-tailed deer.

## Materials and Methods

### White-tailed deer tissues origin and handling

Samples used in this project were opportunistically collected from naturally infected, asymptomatic, CWD-positive white-tailed deer in Hampshire County, West Virginia. The deer were collected during March and April of 2011 and 2012, by West Virginia Division of Natural Resources (WVDNR) during targeted surveillance and management activities (A2018 02-010). Doe CWD status was confirmed by immunohistochemistry (IHC) on medial retropharyngeal lymph node (RPLN) and obex according to standard protocols at the Southeastern Cooperative Wildlife Disease Study (SCWDS; University of Georgia, Athens, GA USA). Fourteen whole white-tailed deer fetuses were collected from nine CWD-positive does, including five sets of twins and four singles. For comparison, control fetal tissues were collected from four CWD-negative deer collected from Georgia, USA, a region where CWD is not known to occur. Shortly after the time of death, adult deer were necropsied in the field and intact fetuses were individually bagged and frozen. Once CWD status of the does was confirmed, frozen fetuses were shipped to Colorado State University (CSU). After thawing, tissues from the fetuses were harvested at CSU using single-use, animal- and tissue-specific blades and forceps to prevent cross-contamination, as previously described ^24^. The following tissues were collected from each fetus: brain, third eyelid, vagus nerve, spleen, colon, ileum, rectum, retropharyngeal lymph node, popliteal lymph node, thymus, liver, and lung. Tissues were then homogenized at 10 or 20% w/v in cold, sterile 0.1 M PBS containing 0.1% Triton-X, using 0.5mm Zirconium Oxide beads in a BBX24B Bullet Blender Blue Homogenizer (NextAdvance). All samples were coded, double-blinded, and then subjected to sPMCA as previously described ^26^. Sample identities were not revealed until after all analyses were completed.

### Gestational age

The fetal gestational age was determined by use of the Hamilton equation; Age (days) = (body length [in mm] x 0.32 + 36.82 ^27^.

### sPMCA

A 10% normal brain homogenate (NBH) in 0.1 M PBS buffer (pH 7.5, with 1% Triton X-100) was prepared from whole brains collected from clinically healthy naïve transgenic mice (<4 months of age) that overexpress PrPcervid (TgCerPrP-E2265037, sourced from Glenn Telling) to serve as a substrate for the PrP^C^ to PrP^CWD^ seeded conversion reaction in sPMCA. Briefly, 30 μl of fetal tissue homogenate was added to 50 μl 10%w/v NBH and subjected to seven 24 hour-rounds of sonication. Each round of sPMCA equals 288 cycles of 10 sec sonication/5 min incubation at 37°C (Misonix). After each round, 30 μl amplified material was transferred to 0.2 ml PCR tubes containing 50 μl 10% NBH, two 2.38 mm and three 3.15 mm Teflon beads (McMaster-Carr). This assay is a highly sensitive and specific method of PrP^res^ detection ^28^, demonstrated to be superior to both IHC and to western blotting of unamplified material ^29^. In order to determine the sensitivity of PMCA in our laboratory, we performed PMCA using serial dilution of known CWD-positive white-tailed deer brain and determined that the sensitivity limit after seven rounds of amplification was reached at 10-11 dilution of 0.2 mg brain 10% homogenate in 0.1 M PBS ^26^. Specificity was ascertained by performing 75 separate reactions, of seven rounds each, using known laboratory negative and positive 10% cervid brain homogenate controls ^26^. We observed no spontaneous amplification in our negative controls, whereas positive controls always had detectable PrP^CWD^ upon proteinase K digestion and visualization by western blotting using an anti-cervid PrP antibody BAR-224 (Cayman Chemical) conjugated to horseradish peroxidase (HRP).

### Western Blotting

The seventh round of sPMCA reaction was assessed by western blotting as previously described ^24,26^. Briefly, known laboratory controls, and unamplified and amplified samples from CWD endemic and non-endemic areas were mixed with proteinase K (Invitrogen) for a 20μg/ml final concentration, and incubated at 37°C for 30 min, followed by an additional 10 min at 45°C with shaking. Samples were mixed with Reducing Agent (10X)/LDS Sample Buffer (4X) (Invitrogen) per manufacturer’s instructions, heated to 95°C for 5 min, then run through a 12% Bis-Tris gel at 100 volts for 2 hr. Proteins were transferred to a polyvinylidene fluoride (PVDF) in a Trans-Blot Turbo Transfer System (BioRad). The membrane was blocked with Casein TBS Blocking buffer (Thermo Scientific) and probed with BAR-224 antibody as described above, then developed with ECL Plus Western Blotting Detection Reagents (GE) and viewed with the ImageQuant LAS-4000 (GE). Sample identities were revealed after blotting.

## Results

To detect the minute amount of amplifiable prions present in fetal tissues, we used sPMCA, a highly sensitive and specific method of PrP^res^ detection ^28^, that is superior to both IHC and western blotting of unamplified material ^29^. Our previous work aimed to assess the presence of CWD prions in fetal tissues harvested from experimentally infected muntjac ^24,25^ and naturally exposed free-ranging elk ^26^. These studies permitted us to optimize assay parameters and the number of sPMCA rounds of amplification necessary to robustly demonstrate the presence of amplification-competent material in each sample ^26^. We therefore deemed it appropriate to examine our naturally sourced samples using these parameters, because we anticipated that vertically transmitted prions, as reported elsewhere would be present in such low concentrations that detection would be difficult without amplification ^24^. Results described here are a subset of the full data analysis. The complete data set will be submitted for peer review.

### Gestational Aging

Using the Hamilton equation ^27^ fetal age ranged between 113 and158 days, with an average age of 133.9 days. Thus, all 18 fetuses (those from CWD+ and age-matched negative control dams) were harvested during the 2^nd^ trimester of pregnancy.

### White-tailed fetal tissue

Prion seeding activity was present in 6 of 14 (43%) fetuses harvested in the 2^nd^ trimester of pregnancy in naturally-infected CWD+ asymptomatic white-tailed deer (Table 1, Figs. 1 and 2).

**Figure 1.**
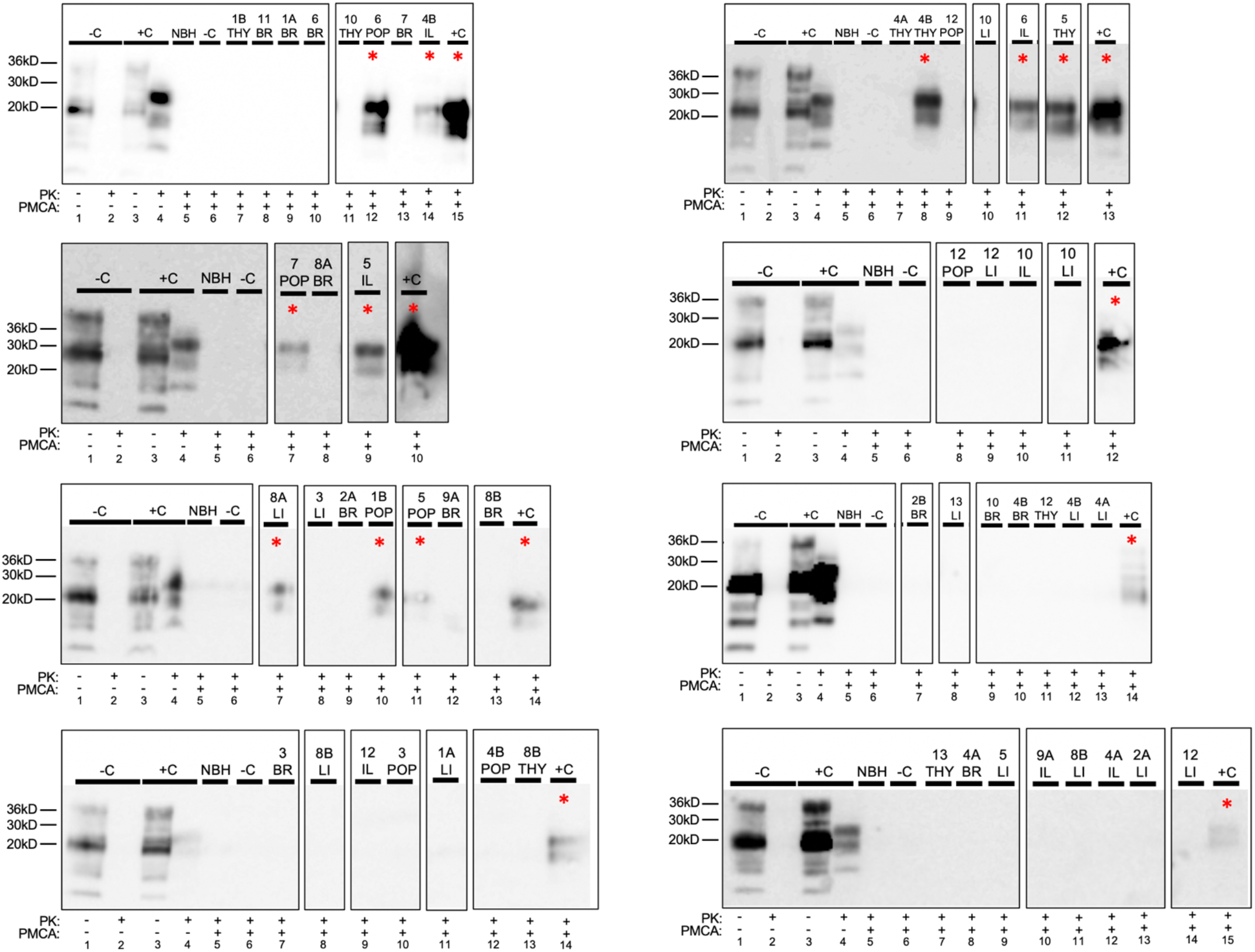
PrP^CWD^ detection in white-tailed deer fetal tissues following seven rounds of sPMCA. Representative western blots for detection of PrP^CWD^ in brain (BR), liver (LI), popliteal lymph node (POP), ileum (IL), and thymus (THY). sPMCA controls (10% homogenate, 7 rounds sPMCA) show complete proteinase K (PK) digestion of negative white-tailed deer brain homogenate (-C) and PK-resistant PrP^CWD^ in CWD-positive brain homogenate (+C). Unamplified western blot assay controls show complete PK-digestion of CWD-negative white-tailed deer brain homogenate (-C;lane 2) and PK-resistant PrP^CWD^ in CWD-positive white-tailed deer brain homogenate (+C;lane 4) (10% homogenate, undiluted, no sPMCA). * = sPMCA positive. NBH = normal brain homogenate. Sample type is identified along the top row of each western blot.

**Fig 2.**
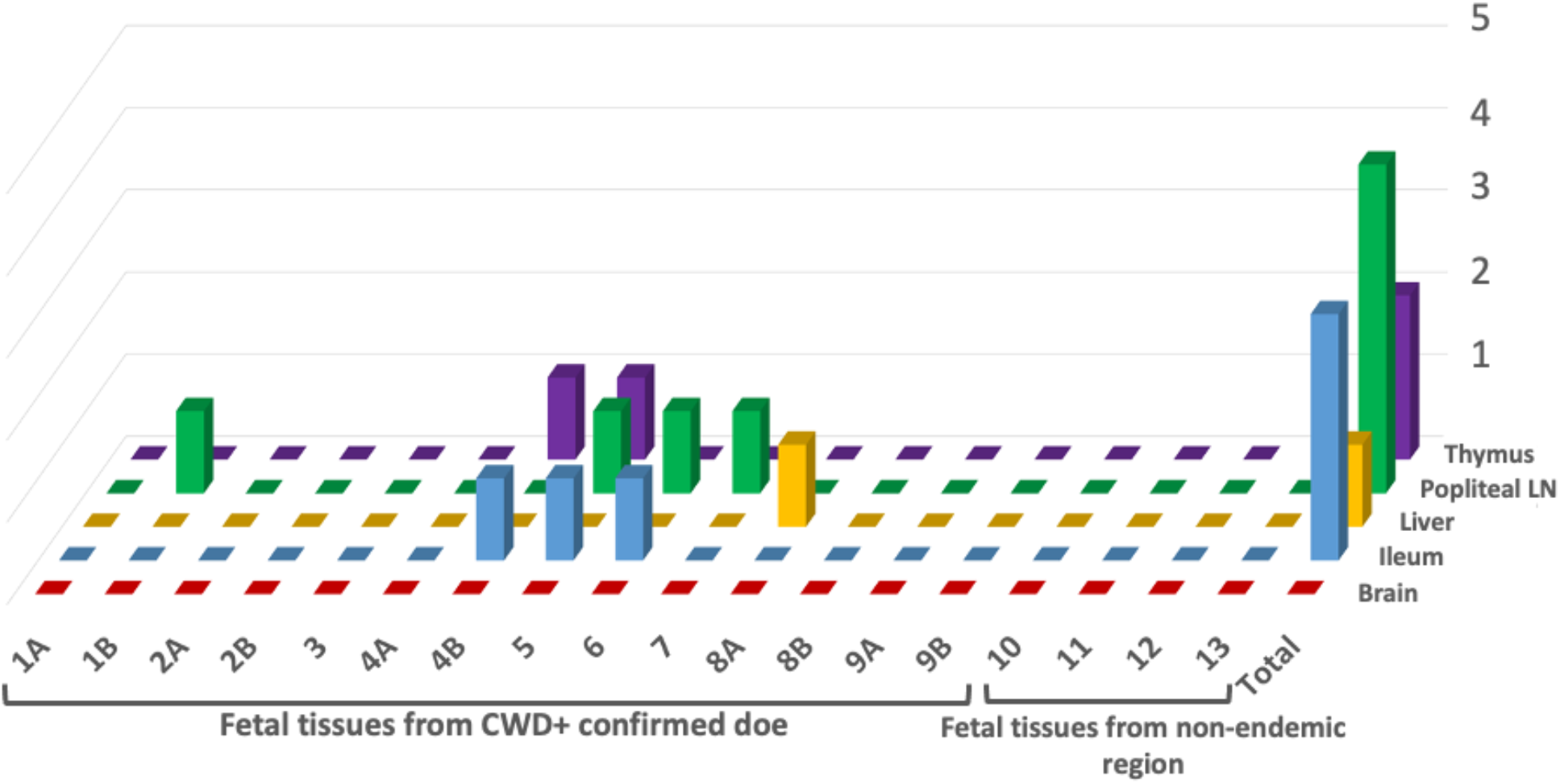
CWD detection in tissues harvested from fetuses of CWD+ white-tailed deer does. All *in utero* derived tissues were analyzed for amplifiable CWD prion seeding activity by 7 rounds of sPMCA. Tissues from fetuses collected in regions non-endemic for CWD were sPMCA negative.

**Table 1.**
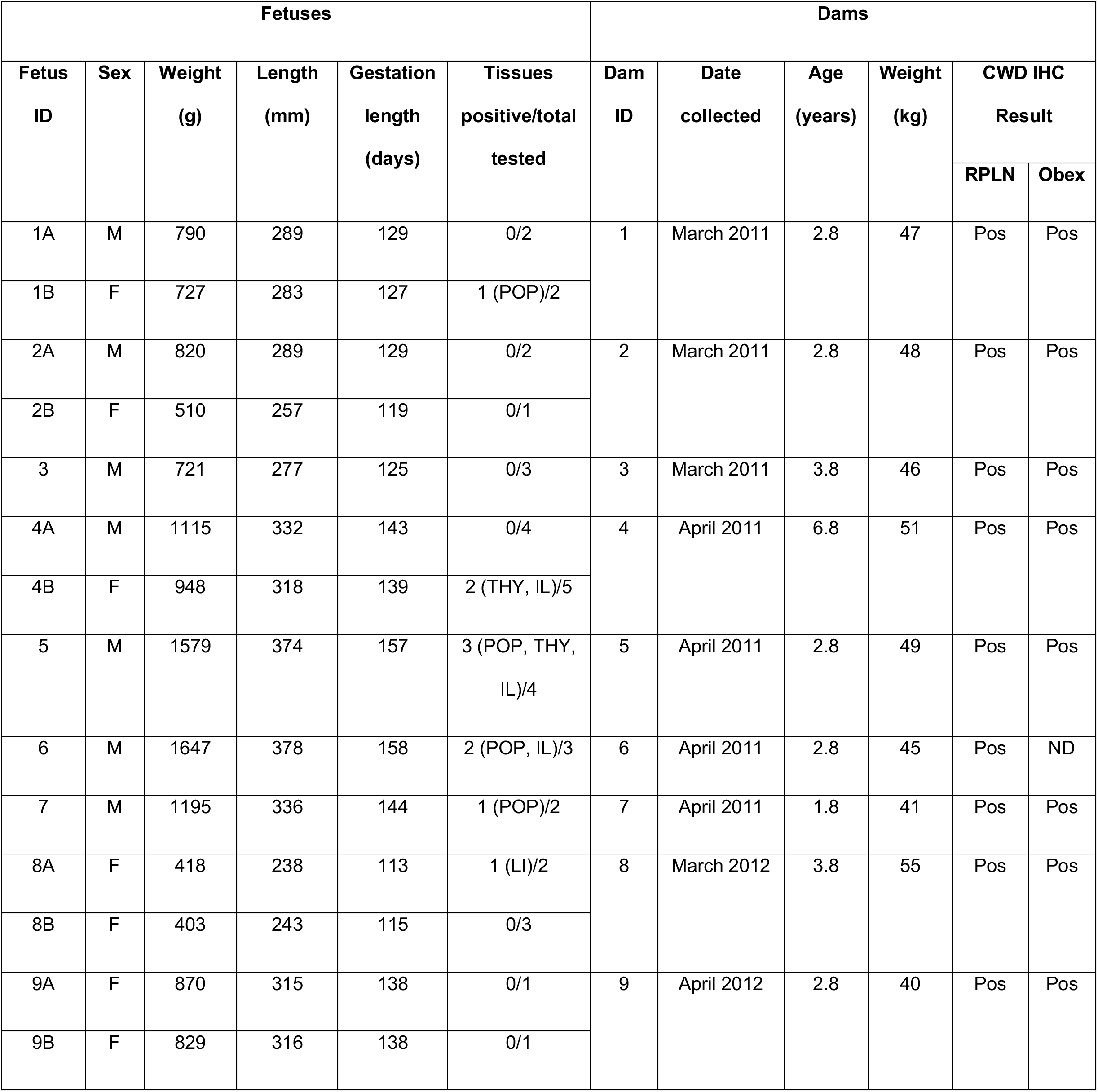
Demographic information from chronic wasting disease-positive presymptomatic, gravid white-tailed deer dams and corresponding fetuses collected in Hampshire County, West Virginia to investigate potential for *in utero* transmission of prions. Tissue codes: POP, popliteal LN; THY, thymus; IL, ileum; LI, liver.

Prion seeding was detected in one of two fetal twins from dams 1, 4 and 8, were positive and neither twin was positive from dams 2 and 9. (Table 1). Fetal tissues containing amplifiable prion seeding activity included liver, popliteal lymph node, ileum, and thymus (Fig. 2). Similar fetal tissues harvested from CWD negative dams remained free of seeding activity after seven rounds of sPMCA (Figs. 1 and 2, Table 1).

## Discussion

The continued geographical expansion of CWD in free-ranging cervid populations prompted us to investigate the role of maternally-derived infections in the facile transmission of this neurodegenerative prion disease. Here we demonstrate prebirth CWD exposure in white-tailed deer. Prion seeds were present in 6 of 14 fetuses collected from the uteri of asymptomatic naturally exposed free-ranging CWD+ white-tailed deer. These results are strikingly similar to our previous findings in free-ranging elk ^26^.

All of the fetuses in this study were opportunistically harvested during the second trimester of pregnancy. Prion seeds were found in an array of fetal tissues, suggesting broad distribution within the fetus during this gestational stage. Studies demonstrating PrP^Scrapie^ deposition in fetal tissues collected from scrapie-infected sheep provided further evidence for prion distribution within fetal tissues of prion-infected dams ^18,30^. We have since demonstrated the presence of prions in early gestational *in utero*-derived fetal tissues of CWD-infected cervids ^24,26^. Further emphasizing that offspring of scrapie and CWD-infected hosts are exposed to prions long before exposure to contaminated environments or postpartum maternal secretions and excretions.

Here we show evidence in three deer for prion accumulation in one of two fetuses in a twin set. Similar findings have been reported in fetal tissues harvested from scrapie-infected sheep ^11^. This suggests the potential for a more focused infection within the pregnancy microenvironment, perhaps within specific placentomes that support fetal growth. The results of our experimental maternal infection studies in muntjac ^25^ and investigation of free-ranging elk ^26^ provide evidence that placentome prion seeding activity can vary within the same pregnancy, with some placentomes containing prions seeds, while other placentomes do not. Studies conducted in the scrapie system have targeted PrP^Scrapie^ deposition within the placentome structure ^11^. With PrP^Scrapie^ accumulation present within all regions of the placentome dependent upon pregnancy stage. Cervid pregnancies are typically supported by 5 to 6 placentomes while sheep pregnancies are supported by 30 or more placentomes. The placentome structure permits the exchange of nutrients and waste between mother and fetus throughout pregnancy ^31^. While the placental structure provides a barrier for the transfer of agents between mother and fetus, leaky or natural breaks within the placental structure are known to occur creating small pools of blood within the maternal-fetal interface ^11^. Blood is known to harbor prion infection ^32–36^. Fetal-derived trophoblast cells are phagocytic and motile ^37–41^. We hypothesize that fetal derived trophoblast cells enter blood-pooled spaces within the placentome, phagocytose prion carrying blood cells and transport them back to the fetus.

Uterine infection would also be an obvious focus of maternal infection that may contribute to fetal infections. Previous findings employing our maternal infection muntjac model, where pregnancy timing and maternal/fetal prion deposition could be closely monitored, suggests that prion seeds are present within the placentome and fetus prior to the uterus ^25^. These findings support the need for further investigation at the maternal-fetal interface seeking mechanisms of prion transfer from mother to offspring.

CWD is the most efficiently transmitted of the prion diseases ^42^. The role of mother to offspring CWD transmission in free-ranging cervid populations remains largely unknown. Understanding the importance of this potential prion transmission route in free-ranging white-tailed deer populations is important to informing control strategies, as well as projecting CWD spread and potential impacts. The findings from this study in free-ranging white-tailed deer, coupled with our previous findings in free-ranging elk, provide the basis for continued exploration of the role vertical transmission may play in cervid populations across North America and other regions of the world, and may broaden our perspective of the transmission dynamics for all prion diseases.

## Acknowledgments

This study was made possible by NIH R01I093634 and the continued financial support from the member states of SCWDS provided by the Federal Aid to Wildlife Restoration Act (50 Stat. 917) and through long-term cooperative agreements with the US Geological Survey Ecosystems Management Area, and US Department of Agriculture, Animal Plant Health Inspection Service. We thank David Osborn for nonendemic samples, and Brandon Munk and John Wlodkowski (SCWDS) for laboratory assistance.

## Notes

### Competing Interest Statement

The authors have declared no competing interest.

## References

1 Mathiason, C. K. et al. Infectious prions in the saliva and blood of deer with chronic wasting disease. Science 314, 133–136, doi:10.1126/science.1132661 (2006).

2 Tamguney, G. et al. Asymptomatic deer excrete infectious prions in faeces. Nature 461, 529–532, doi:10.1038/nature08289 (2009).

3 Safar, J. G. et al. Transmission and detection of prions in feces. The Journal of infectious diseases 198, 81–89, doi:10.1086/588193 (2008).

4 Haley, N. J., Seelig, D. M., Zabel, M. D., Telling, G. C. & Hoover, E. A. Detection of CWD prions in urine and saliva of deer by transgenic mouse bioassay. PloS one 4, e4848, doi:10.1371/journal.pone.0004848 (2009).

5 Hoinville, L. J., Tongue, S. C. & Wilesmith, J. W. Evidence for maternal transmission of scrapie in naturally affected flocks. Prev Vet Med 93, 121–128, doi:10.1016/j.prevetmed.2009.10.013 (2010).

6 Dickinson, A. G., Young, G. B., Stamp, J. T. & Renwick, C. C. An analysis of natural scrapie in Suffolk sheep. Heredity 20, 485–503 (1965).

7 Dickinson AG, S. J., Renwich CC. Maternal and lateral transmission of scrapie in sheep. J Comp Pathol 84, 19–25 (1974).

8 Andreoletti O, L. C., Chabert A, Monnereau L, Tabouret G, Lantier F, Berthon P, Eychenne F, Lafond, Benestad S, Elsen JM, Schelcher F. PrPSc accumulation in placentas of ewes exposed to natural scrapie: influence of foetal PrP genotype and effect on ewe-to-lamb transmission. 83, 2607–2616 (2002).

9 O’Rourke, K. I., Zhuang, D., Truscott, T. C., Yan, H. & Schneider, D. A. Sparse PrP(Sc) accumulation in the placentas of goats with naturally acquired scrapie. BMC veterinary research 7, 7, doi:10.1186/1746-6148-7-7 (2011).

10 Tuo, W. et al. Prp-c and Prp-Sc at the fetal-maternal interface. The Journal of biological chemistry 276, 18229–18234, doi:10.1074/jbc.M008887200 (2001).

11 Tuo, W. et al. Pregnancy status and fetal prion genetics determine PrPSc accumulation in placentomes of scrapie-infected sheep. Proceedings of the National Academy of Sciences of the United States of America 99, 6310–6315, doi:10.1073/pnas.072071199 (2002).

12 Onodera, T., Ikeda, T., Muramatsu, Y. & Shinagawa, M. Isolation of scrapie agent from the placenta of sheep with natural scrapie in Japan. Microbiology and immunology 37, 311–316 (1993).

13 Lacroux, C. et al. Dynamics and genetics of PrPSc placental accumulation in sheep. The Journal of general virology 88, 1056–1061, doi:10.1099/vir.0.82218-0 (2007).

14 Foster, J. D., Goldmann, W. & Hunter, N. Evidence in sheep for pre-natal transmission of scrapie to lambs from infected mothers. PloS one 8, e79433, doi:10.1371/journal.pone.0079433 (2013).

15 Alverson, J., O’Rourke, K. I. & Baszler, T. V. PrPSc accumulation in fetal cotyledons of scrapie-resistant lambs is influenced by fetus location in the uterus. The Journal of general virology 87, 1035–1041, doi:10.1099/vir.0.81418-0 (2006).

16 Race, R., Jenny, A. & Sutton, D. Scrapie infectivity and proteinase K-resistant prion protein in sheep placenta, brain, spleen, and lymph node: implications for transmission and antemortem diagnosis. The Journal of infectious diseases 178, 949–953 (1998).

17 Schneider, D. A. et al. The placenta shed from goats with classical scrapie is infectious to goat kids and lambs. The Journal of general virology 96, 2464–2469, doi:10.1099/vir.0.000151 (2015).

18 Spiropoulos, J., Hawkins, S. A., Simmons, M. M. & Bellworthy, S. J. Evidence of in utero transmission of classical scrapie in sheep. J Virol 88, 4591–4594, doi:10.1128/JVI.03264-13 (2014).

19 Maddison, B. C. et al. Prions are secreted in milk from clinically normal scrapie-exposed sheep. J Virol 83, 8293–8296, doi:10.1128/JVI.00051-09 (2009).

20 Konold, T., Moore, S. J., Bellworthy, S. J. & Simmons, H. A. Evidence of scrapie transmission via milk. BMC veterinary research 4, 14, doi:10.1186/1746-6148-4-14 (2008).

21 Konold, T. et al. Evidence of effective scrapie transmission via colostrum and milk in sheep. BMC veterinary research 9, 99, doi:10.1186/1746-6148-9-99 (2013).

22 Madsen-Bouterse, S. A., Highland, M. A., Dassanayake, R. P., Zhuang, D. & Schneider, D. A. Low-volume goat milk transmission of classical scrapie to lambs and goat kids. PloS one 13, e0204281, doi:10.1371/journal.pone.0204281 (2018).

23 Ligios, C. et al. Sheep with scrapie and mastitis transmit infectious prions through the milk. J Virol 85, 1136–1139, doi:10.1128/JVI.02022-10 (2011).

24 Nalls, A. V. et al. Mother to offspring transmission of chronic wasting disease in reeves’ muntjac deer. PloS one 8, e71844, doi:10.1371/journal.pone.0071844 (2013).

25 Nalls, A. V. et al. Infectious Prions in the Pregnancy Microenvironment of CWD-infected Reeves’ Muntjac Deer. J Virol pii: JVI.00501-17, doi:10.1128/JVI.00501-17 (2017).

26 Selariu, A. et al. In utero transmission and tissue distribution of chronic wasting disease-associated prions in free-ranging Rocky Mountain elk. The Journal of general virology, doi:10.1099/jgv.0.000281 (2015).

27 Hamilton RJ, T. M., Moore WG. in Proc Annu Conf Southeast Assoc Fish Wildl Agencies 389e395.

28 Saa, P., Castilla, J. & Soto, C. Presymptomatic detection of prions in blood. Science 313, 92–94, doi:10.1126/science.1129051 (2006).

29 McNulty, E. et al. Comparison of conventional, amplification and bio-assay detection methods for a chronic wasting disease inoculum pool. PloS one 14, e0216621, doi:10.1371/journal.pone.0216621 (2019).

30 Garza MC, F.-B. N., Boles R, Badiola JJ, Castilla J, Monleon E. Detection of PrPres in Genetically Susceptible Fetuses from Sheep with Natural Scrapie. PloS one 6, e27525, doi:10.1371/journal.pone.0027525.g001 (2011).

31 Slack, J. M. W. Essential Developmental Biology. (2013).

32 Hunter, N. et al. Transmission of prion diseases by blood transfusion. The Journal of general virology 83, 2897–2905, doi:10.1099/0022-1317-83-11-2897 (2002).

33 Houston, F., Foster, J. D., Chong, A., Hunter, N. & Bostock, C. J. Transmission of BSE by blood transfusion in sheep. Lancet 356, 999–1000 (2000).

34 Mathiason, C. K., Powers, J. G. & Dahmes, S. J. Don’t kiss that deer. Compendium on Continuing Education For the Practicing Veterinarian 28, 822 (2006).

35 Mathiason, C. K. et al. B Cells and Platelets Harbor Prion Infectivity in the Blood of Deer Infected with Chronic Wasting Disease. Journal of Virology 84, 5097–5107, doi:10.1128/JVI.02169-09 (2010).

36 Peden, A. H., Head, M. W., Ritchie, D. L., Bell, J. E. & Ironside, J. W. Preclinical vCJD after blood transfusion in a PRNP codon 129 heterozygous patient. Lancet 364, 527–529, doi:10.1016/S0140-6736(04)16811-6 (2004).

37 Myagkaya, G. & Schellens, J. P. Final stages of erythrophagocytosis in the sheep placenta. Cell and tissue research 214, 501–518 (1981).

38 Wooding, F. B. The role of the binucleate cell in ruminant placental structure. J Reprod Fertil Suppl 31, 31–39 (1982).

39 Wooding, F. B. Current topic: the synepitheliochorial placenta of ruminants; binculeate cell fusions and hormone production. Placenta 13, 101–113 (1992).

40 Wooding, F. B. Marshall’s Physiology of Reproduction. 4th edn, Chapter 4 (London, 1994).

41 Wooding, F. B., Morgan, G., Adam, C.L. Structure and function in the rumiant synepithelialchorial placenta: central role of the trophoblast binucleate cll in deer. Microsc Res Tech 38, 88–99 (1997).

42 Benestad, S. L. & Telling, G. C. Chronic wasting disease: an evolving prion disease of cervids. Handb Clin Neurol 153, 135–151, doi:10.1016/B978-0-444-63945-5.00008-8 (2018).

